# A CNN model for predicting binding affinity changes between SARS-CoV-2 spike RBD variants and ACE2 homologues

**DOI:** 10.1101/2022.03.22.485413

**Authors:** Chen Chen, Veda Sheersh Boorla, Ratul Chowdhury, Ruth H. Nissly, Abhinay Gontu, Shubhada K. Chothe, Lindsey LaBella, Padmaja Jakka, Santhamani Ramasamy, Kurt J. Vandegrift, Meera Surendran Nair, Suresh V. Kuchipudi, Costas D. Maranas

**Author notes:** Author contributions: C.C., V.S.B., C.D.M., S.V.K. designed research; C.C., V.S.B. performed research; C.C., V.S.B. analyzed data; C.C., V.S.B., C.D.M., R.C., S.V.K., S.K.C, K.J.V., R.H.N., A.G., M.S.N., L.L., P.J., S.R. wrote the paper. C.C. and V.S.B. contributed equally to this work.

## Abstract

The cellular entry of severe acute respiratory syndrome coronavirus 2 (SARS-CoV-2) involves the association of its receptor binding domain (RBD) with human angiotensin converting enzyme 2 (hACE2) as the first crucial step. Efficient and reliable prediction of RBD-hACE2 binding affinity changes upon amino acid substitutions can be valuable for public health surveillance and monitoring potential spillover and adaptation into non-human species. Here, we introduce a convolutional neural network (CNN) model trained on protein sequence and structural features to predict experimental RBD-hACE2 binding affinities of 8,440 variants upon single and multiple amino acid substitutions in the RBD or ACE2. The model achieves a classification accuracy of 83.28% and a Pearson correlation coefficient of 0.85 between predicted and experimentally calculated binding affinities in five-fold cross-validation tests and predicts improved binding affinity for most circulating variants. We pro-actively used the CNN model to exhaustively screen for novel RBD variants with combinations of up to four single amino acid substitutions and suggested candidates with the highest improvements in RBD-ACE2 binding affinity for human and animal ACE2 receptors. We found that the binding affinity of RBD variants against animal ACE2s follows similar trends as those against human ACE2. White-tailed deer ACE2 binds to RBD almost as tightly as human ACE2 while cattle, pig, and chicken ACE2s bind weakly. The model allows testing whether adaptation of the virus for increased binding with other animals would cause concomitant increases in binding with hACE2 or decreased fitness due to adaptation to other hosts.

## 1. Introduction

The severe acute respiratory syndrome coronavirus 2 (SARS-CoV-2) is responsible for the coronavirus disease 2019 (COVID-19) pandemic and has continued to evolve since the initial outbreak.^1^ Several variants of the wild-type (WT) virus (Wuhan-Hu-1)^2^ have been identified in different countries, including the United Kingdom (Alpha or B.1.1.7, Eta or B. 1.525), South Africa (Beta or B.1.351, Omicron or B.1.1.529 or BA.1), Brazil (Gamma or P.1, Zeta or P.2), United States (Epsilon or B.1.429/B.1.427, Iota or B.1.526), India (Kappa or B.1.617.1, Delta or B.1.617.2), Philippines (Theta or P.3), Columbia (Mu or B.1.621), Peru (Lambda or C.37) and France (B.1.640.2).^3–5^ The Omicron variant has quickly overtaken Delta as the globally dominant variant thanks in part to 60 mutations which have conferred increased transmissibility and increased reinfection risk as well as significant reductions in vaccine effectiveness.^6,7^ The Omicron subvariant BA.2 contains additional amino acid changes, and it has been reported that both naive and immunologically trained individuals exhibit even higher susceptibility to BA.2 than the original Omicron strain.^8^ While novel therapeutics and authorized COVID-19 vaccines have been deployed to mitigate the public health impacts of SARS-CoV-2 and limit its spread, the continuous emergence of SARS-CoV-2 variants is complicating the situation and results in periodic surges of new infections.^9^

Many domestic and wild animals have been shown to be susceptible to SARS-CoV-2 either by natural and/or experimental infections,^9,10^ revealing an extensive host range of SARS-CoV-2. Cats, dogs, lions, tigers in zoos, minks, and ferrets have been reported to be infected via contact with COVID-19 patients, while snow leopards, pumas, and gorillas have been found to be infected with SARS-CoV-2 by natural means.^10^ White-tailed deer have also been confirmed to be susceptible to SARS-CoV-2 through infection studies^11,12^ After anti-SARS-CoV-2 antibodies were detected in 33% of the 481 white-tailed deer samples in four different states^13^, multiple reports of natural infection of deer in USA^14–17^ and Canada^18,19^ have been reported. Moreover, recent studies^20^ showed that Syrian hamsters in Hong Kong and deer in Canada^18^ were able to transmit SARS-CoV-2 to humans, adding to the list of animal to human transmissions after the initial documented case in mink^21^ with evidence for animal-to-human spillover. These findings underscore the importance of continual monitoring of potential spillovers of SARS-CoV-2 into non-human species, as the virus could gain a toe-hold and evolve in animal reservoirs, which could constitute a persistent threat of spillover back to humans. This scenario would complicate future mitigation strategies against the SARS-CoV-2 virus.^21^

SARS-CoV-2 is an enveloped single-stranded RNA virus that expresses the spike protein on its surface, which mediates the binding to host cells.^22^ The association of the receptor binding domain (RBD) of the spike protein with the human angiotensin converting enzyme-2 (hACE2) receptor represents the first crucial step of viral infection.^2,23^ SARS-CoV-2 variants contain single or multiple amino acid substitution as well as indels in the spike protein, and some of these changes have been shown to alter the RBD-hACE2 binding strength.^3,4^ Several amino acid changes in the RBD of spike protein have considerably impacted the transmissibility and antigenicity in circulating variants, and the increased frequency of these amino acid changes may indicate a positive association with RBD-hACE2 binding affinity enhancement.^3,24^ The amino acid change D614G showed increased prevalence which emerged multiple times in the global SARS-CoV-2 population, and the advantages for infectivity and transmissibility of this mutation have been indicated in several studies.^25–28^ The N439K mutation was observed with increased frequency when circulating widely in Europe,^29^ and Y453F was detected in Denmark among farmed mink and humans,^21^ where both mutations showed enhanced RBD binding affinity to hACE2. Being present in five circulating variants, N501Y can significantly increase the binding affinity by introducing new aromatic stacking interactions and hydrogen bonds.^30,31^ E484K is even more widely prevalent and is present in seven circulating variants. It destabilizes the RBD-down state to favor the RBD-up state which is a required conformation for effective binding to hACE2.^32,33^ In addition to affinity enhancement, E484K also affords escape from neutralizing antibodies,^33–36^ and the presence of the alternative change E484Q in the Kappa variant further suggests the importance of monitoring amino acid changes at this position. In the Omicron variant, as many as 60 amino acid changes are encoded including 37 in the spike protein (with 15 in the RBD),^6^ among which N501Y, T478K and K417N were also identified in earlier variants Alpha, Beta, Gamma, Delta, Theta, and Mu.^3^ The immune escape capabilities are possibly conferred by K417N,^37^ S477N,^38^ E484A^38^ and other novel mutations, while the binding affinity between RBD and hACE2 is retained due to strongly binding-improving mutations such as N501Y,^30,31^ both contributing to the increased viral transmissibility of Omicron compared to earlier variants.^39–41^ Early on during the pandemic, to gain a more complete view of the effect of RBD mutation, Starr et al.^30^ exhaustively assessed the impact of single amino acid changes on RBD expression and hACE2 binding. They found that 84.5% of the amino acid changes are detrimental, 7.5% are neutral and the rest 8.0% can lead to enhanced binding of the RBD with hACE2. Chan et al.^42^ used deep mutagenesis to systematically evaluate the binding affinity of WT RBD for hACE2 mutants, and the engineered decoy receptor for SARS-CoV-2 showed comparably high affinity to neutralizing antibodies. These findings highlight the importance of monitoring single amino acid changes and their potential for binding affinity improvement and call for prospective methods that can rapidly scan and identify amino acid change combinations that are likely to further boost affinity with ACE2.

By pre-screening mutations in search of potentially contagious variants, computational methods can contribute to understanding alterations in the characteristics of the circulating variants and identify particularly problematic ones. The binding free energy can be reasonably approximated using molecular-mechanics-based empirical force fields such as Rosetta.^43–45^ It can also be calculated by performing molecular mechanics-generalized Born surface area (MM-GBSA)^46–49^ analysis on configurations generated by molecular dynamics (MD) simulations.^50^ The hybrid quantum mechanics/molecular mechanics (QM/MM) approaches^51^ can also be used when critical chemical reactions are involved in binding activities, but the significant computational cost restricts QM treatment for large protein-protein complexes.^52^ Reweighting of energy terms can be used to reach better prediction accuracy, where the weights are trained to accurately reproduce experimentally determined binding free energies (ΔΔ*G*_bind_).^53,54^ As a powerful complement, machine learning algorithms can effectively “learn” highly complex relationships among energy components and the target binding affinity from training samples.^55,56^ Using a data-driven approach, higher-level correlations can be captured and later be used to improve the accuracy of predictions.^57^

Recently, we developed a two-step framework to quantitatively predict binding affinity change upon amino acid substitutions.^58^ The first step consists of 48 parallel 4-ns MD simulations of each RBD variant complexed with hACE2, followed by MM-GBSA analysis to extract decomposed binding energy terms. The second step involves implementing a neural network (NN) to predict the apparent dissociation constant (*K*_D,app_) ratios between the variants and the WT using the obtained decomposed energy terms as descriptors. Agreement between experimental values and the NN_MM-GBSA model predictions was significantly better than predictions made directly using raw energies, reaching a correlation coefficient of 0.73 and an accuracy of 82.8% for the classification of improving or worsening the binding affinity.^58^ Albeit encouraging, the computational demand in dataset preparation involving expensive MD simulations prevents the NN_MM-GBSA model from being expanded to a much larger training set and/or from being extensively used for large-scale genomic screening. Therefore, developing a tool with comparable accuracy yet better computational efficiency remains relevant. Readily accessible descriptors are crucial for machine learning processes, and the protein sequence is a promising candidate. Recent breakthroughs by AlphaFold2^59^ and RoseTTAFold^60^ demonstrated that the amino acid sequence contains a wealth of information for protein structure prediction, which can serve as the basis for protein-protein binding affinity prediction.^61–65^

Herein, we introduce a convolutional neural network (CNN) regression model (CNN_seq) based on protein sequence and WT complex structural features. Various features for (i) individual residue identities including hydropathy index,^66^ volume,^67^ zScales,^57^ and VHSE^68^ (principal components score Vectors of Hydrophobic, Steric, and Electronic properties), (ii) residue-pair interactions in AAIndex^69^, and (iii) residue properties such as normalized vdW volume, polarity, charge, polarizability, secondary structure, and solvent accessibility^69^ were used in the feature encoding procedure. The training was performed against all 8,440 variants with single and multiple amino acid changes in the RBD and hACE2. The predictive capability of this CNN_seq model was assessed by comparing it with the experimental *K*_D,app_ ratios, achieving a classification accuracy of 83.28% and a correlation coefficient *r* of 0.85 in five-fold cross-validation tests. The model was further tested against many circulating variants that were blind to the model during training, and most were predicted to show improved binding affinity, as demonstrated by other experimental studies.^23–25,29,30,32,37,70–91^ Furthermore, we randomly chose 1,667 of the 8,440 variants to serve as the blind test set and used the rest to perform a fivefold cross-validation test. Similar performance (%VC = 83.47%, *r* = 0.84) was observed for the blind test set, reconfirming the robustness of the CNN model predictions.

Given the accuracy and efficiency of this CNN_seq model, we were able to exhaustively screen over 220,000 RBD variants for each host with combinations of up to four amino acid substitutions and suggest candidates with the highest improvement in RBD-ACE2 binding affinity for monitoring. The predicted binding affinity of RBD variants for deer ACE2 was found to be similar to humans, which was lower for cattle and pigs and lowest for chickens. The computational model can be accessed from the GitHub repository (https://github.com/maranasgroup/CNN_seq_CoV2).

## 2. Results

### Dataset preparation

We curated the training dataset by combining experimental results of deep mutational scanning of RBD (Starr et al.^30^) and random mutational scanning of hACE2 (Chan et al.^42^). The curated dataset consisted of 8,440 variants (see Methods for details of variants selection and distribution). To quantify the change in RBD binding affinity for hACE2, Starr et al.^30^ reported the apparent dissociation constant *K*D,app ratio which was defined as *K*_D,app,variant_/*K*_D,app,WT_. For each variant, as compared to the WT RBD, a *K*_D,app_ ratio greater than 1 indicates stronger (or improving) binding to hACE2 whereas a value less than 1 implies weaker (or worsening) binding. Note that, the *K*_D,app,WT_ in present work always refers to the *K*_D,app_ for the complex formed by WT hACE2 and WT SARS-CoV-2 RBD proteins, and such designation allows for the direct comparison across human and animal hosts.

### Feature encoding

The CNN_seq model uses both sequence-based and structure-based features in model construction to maximize the utilization of available data. For the human case, the sequences and three-dimensional (3D) structure of SARS-CoV-2 spike RBD in complex with hACE2 were obtained from the Protein Data Bank^92^ (PDB ID: 6LZG), for the animal cases, the sequences of ACE2 proteins were collected from UniProt^93^ for four species: deer (*Odocoileus virginianus*, ID: A0A6J0Z472), cattle (*Bos indicus* × *Bos taurus*, ID: A0A4W2H6E0), pig (*Sus scrofa*, ID: A0A220QT48), and chicken (*Gallus gallus*, ID: F1NHR4). Due to the lack of experimentally determined structures for these animals, we used SWISS-Model^94^ to perform homology modeling and generated 3D structures for the complexes. Subsequent structural refinement for each complex was performed through an MD simulation with explicit water, and the interface equilibration was demonstrated by reaching a standard deviation (SD) in the root mean square deviation (RMSD) of less than 1 Å in the final 100-ns (see Methods for details).

The structure of the RBD-ACE2 complex encodes information as to the local environment that is accessible by each residue, and the interactions between residues in ACE2 and RBD underpin the overall binding strength of the complex. We used the PISA (Proteins, Interfaces, Structures and Assemblies) tool^95^ provided by Protein Data Bank in Europe (PDBe)^96^ to help identify the interfacial residues as well as key interactions such as hydrogen bonds, salt bridges, and disulfide bonds formed by compatible residue pairs (Figure 1A). In addition, the Protein Contact Maps tool developed by Benjamin et al.^97^ was employed to generate the contact map (Figure 1B), which is a two-dimensional (2D) matrix recording the distances between all possible amino acid residue pairs in the complex. For each complex, the structure, contact map and the interfacial residue information are not changed significantly when mutations are subsequently introduced.

**Figure 1.**
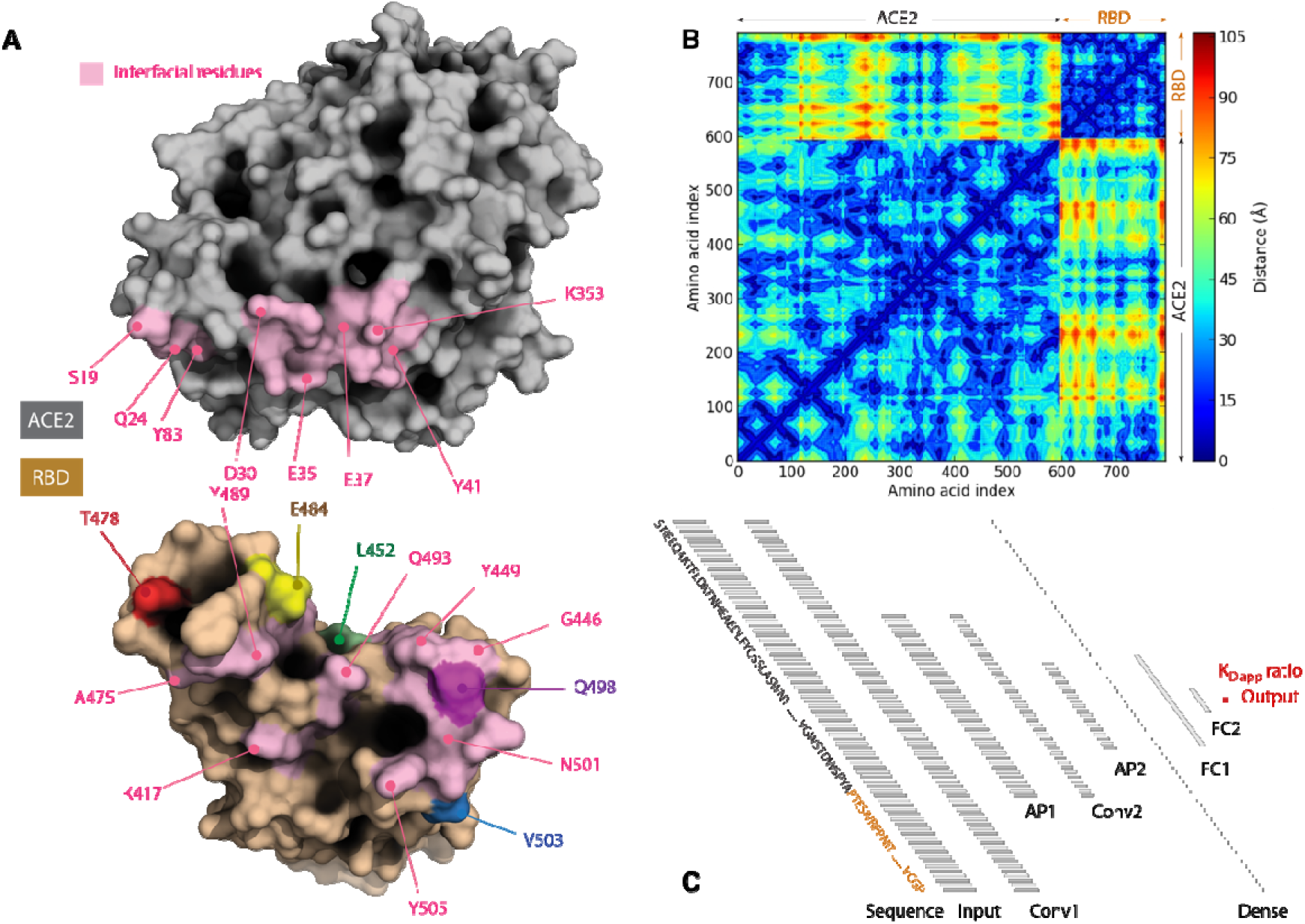
Schematic representation of the CNN_seq workflow. (A) Identification of interfacial residues, hydrogen bonds, salt bridges and disulfide bonds from the structure of receptor binding domain (RBD) complexed with angiotensin converting enzyme-2 (ACE2). Interfacial residues are colored pink on protein surfaces, and some key non-interfacial residues are highlighted. (B) Generation of the distance map recording the distances between all residue pairs in the complex structure. (C) The structure of CNN_seq model which takes RBD-ACE2 complex sequence as the input and apparent dissociation constant (*K*_D,app_) ratio as the output.

### CNN model trained on experimental *K*_D,app_ ratio

A CNN regression model was constructed by taking the RBD-ACE2 sequence for variants as the input. As illustrated in Figure 1C, the CNN_seq model contains one input layer, two 1D convolutional (Conv) layers, two 1D average pooling (AP) layers, one dense layer, two fully-connected (FC) layers, and an output layer. There is a rectified linear unit (ReLU) activation function following each Conv and FC layer, and dropout regularization method^98^ is applied to each FC layer to reduce overfitting (see Methods for details of the CNN structure). The matrix of input features pass through a series of layers and functions continuously, and the final output is the apparent dissociation constant *K*_D,app_ ratio. Such architecture takes into account the interactions between individual residues and their local environment, and allows higher level correlations between domains to be captured.

For model evaluation, we followed the five-fold cross-validation procedure, where the entire dataset was split into five subsets, with each subset considered as the test set once, and the rest of the four subsets are used together to form the training set. A complete cycle of five-fold crossvalidation produced five individual models, and each model made predictions on the subset of variants that the model itself did not see. Combining predictions of the five subsets, all the variants in the original dataset were predicted once, and the performance of the model can be fairly evaluated. Two metrics were used to evaluate the performance of the CNN_seq model, the percent recovery of correct variant classification (%VC) and the Pearson correlation coefficient (*r*). The %VC calculates in percentage the accuracy of classifying the direction of change in the binding affinity compared to WT, while *r* measures the strength of the linear correlation between the predicted and experimental *K*_D,app_ ratio values (see Methods for details).

The averaged results from the five-fold cross-validation tests of the CNN_seq model achieved a %VC of 83.28% and a correlation coefficient *r* of 0.85 for a complete dataset of 8,440 variants. Compared to the performance of the NN_MM-GBSA model (%VC = 82.8%, *r* = 0.73),^58^ the %VC of the CNN_seq model remained similar whereas there was a slight improvement in the correlation coefficient *r*. It appears that on balance the information content embedded in the 8,440 variants dataset approximately matches the information contained in the energy terms of the GBSA molecular dynamic simulation of the 108 variants used in model NN_MM-GBSA.^58^ Figure 2 compares the *K*_D,app_ ratio values between experiments and CNN_seq predictions, where 54.94% of the incorrectly classified variants appear to be for variants with experimental *K*_D,app_ ratio values within [0.9-1.1]. The performance of the CNN_seq model was also evaluated separately on ACE2 and RBD variants (Figure S1 in *SI Appendix*). We found that CNN performed better for the 6,105 RBD variants (i.e., %VC = 85.2% and *r* = 0.87) compared to the 2,335 hACE2 variants (%VC = 72.19% and *r* = 0.63). The apparent difference in the performance of CNN in the two datasets alludes to systematic differences in the way affinities were calculated or different error margins. However, since there are nearly three times more RBD variants (72.10% of the dataset) than ACE2 variants (27.90% of the dataset), it is not surprising to see that the performance bias leans towards RBD variants. Although a model built on solely RBD variants could achieve better overall performance, we chose to include both sets of data in the training set because this model was ultimately designed to predict the binding affinity changes for animal hosts, whose ACE2s differ slightly from the human version.

**Figure 2.**
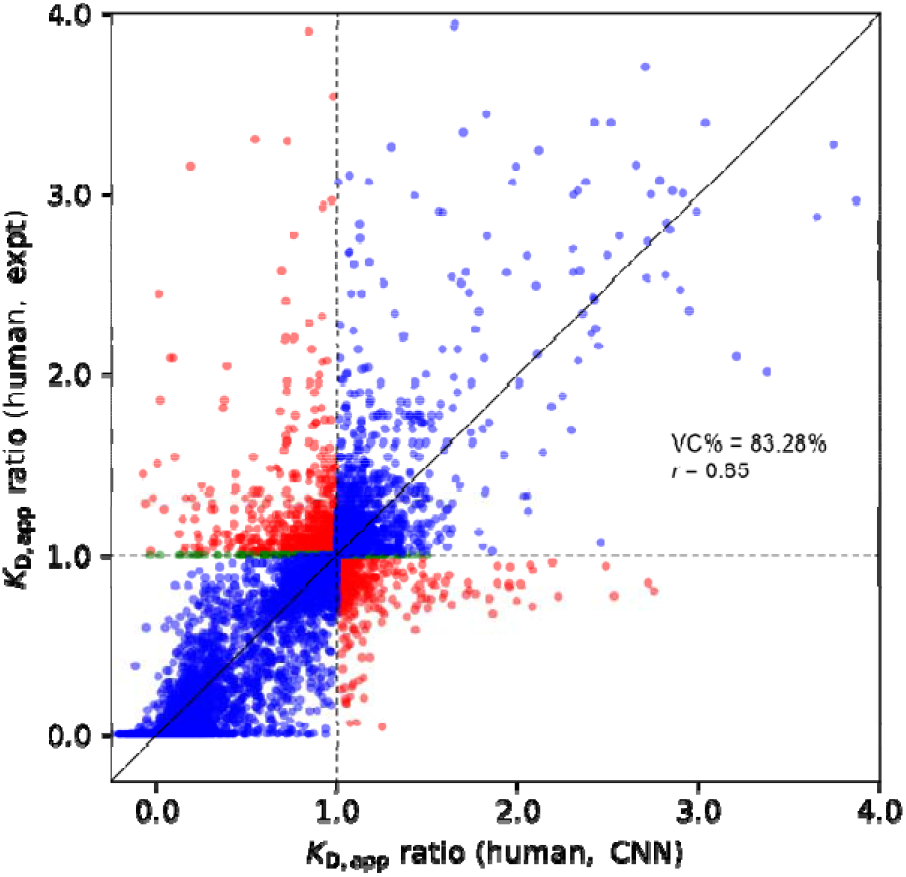
Comparison of *K*_D,app_ ratio between experiments and CNN_seq models predictions from five-fold cross-validation tests on 8,440 variants. Correctly classified variants are colored in blue, incorrectly classified variants are colored in red, and variants with unchanged binding affinities are colored in green. Horizontal and vertical dashed lines are drawn to indicate the dividing line where *K*_D,app_ ratio equals to 1, and a diagonal solid line is drawn to indicate perfect correlation.

### CNN_seq model prediction of *K*_D,app_ of circulating SARS-CoV-2 variants

The circulating variants include amino acid substitutions and/or indels in the SARS-CoV-2 spike protein such as the amino-terminal domain, the RBD, and the furin cleavage sequence.^24^ In total, spike variants for 21 circulating variants were chosen to be examined by the CNN_seq model, including 13 that are assigned WHO labels^99^ and eight that only have Pango^100^ classifications. We also assessed different instances of variants containing amino acid changes only present in some sequences. The experimental values were collected from available experimental results obtained through surface plasmon resonance,^29,32,70,71,89–91^ bio-layer interferometry,^23,25,29,30,37,72–85,90^ enzyme-linked immunosorbent assay,^74,82,86,87,90^ or microscale thermophoresis.^88^ For all variants with single amino acid changes, in the spike protein, we defaulted to the *K*_D,app,variant_/*K*_D,app,WT_ value from the work of Starr et al.^30^ For experimentally tested variants with multiple amino acid changes we used the reported *K*_D,variant_/*K*_D,WT_ to make comparisons. Given the often-large differences in the reported values, the median calculated from the available experiment results was used.

As shown in Table 1, both the CNN_seq and NN_MM-GBSA models predicted improved binding affinity for most circulating variants relative to the WT, consistent with experimental observations. To further assess the performance of CNN_seq and NN_MM-GBSA models on blinded datasets, we also prepared a list of 15 variants containing multiple amino acid changes (Table S1 in *SI Appendix*). For this small blinded test set, the CNN_seq model achieved a %VC of 92.9% and *r* of 0.60 whereas NN_MM-GBSA model performance was somewhat worse with %VC of 75.7% and *r* of 0.28. This performance gap could be because NN_MM-GBSA model was trained using only 108 single amino acid change variants, whereas the blinded dataset involved single amino acid changes with 66.67% not present in the dataset used for training NN_MM-GBSA. Roughly, all circulating variants can be separated into two families, ones with *K*_D,app_ ratios above 1.5 including Alpha, Beta, Gamma, Theta, Mu, B.1.640.2, C.1.2 variants, and the others with *K*_D,app_ ratios between 1 and 1.5 including Delta, Epsilon, Zeta, Eta, Iota, Kappa, Lambda, Omicron, B.1.1.298, B.1.1.519, B.1.1.36, B.1.1.317, B.1.1.141, BA.2. This classification of the variants agrees well with the antigenic map,^1^ alluding to the partial correlation between RBD-ACE2 binding affinity and evolution trajectories.

**Table 1.**
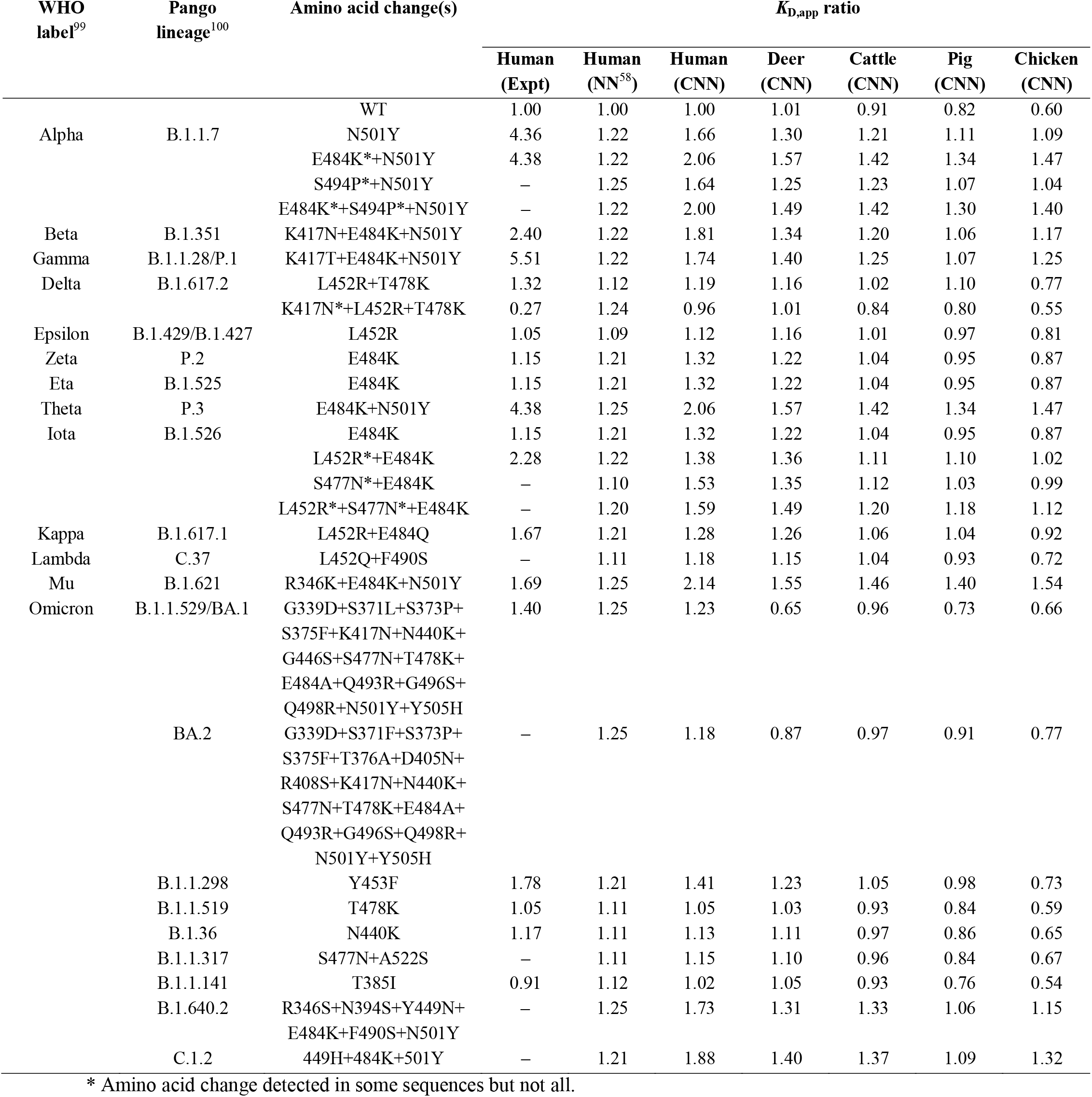
Comparison of *K*_D,app_ ratio from experiments (Expt), CNN_seq (CNN), and NN_MM-GBSA (NN) model predictions on circulating RBD variants complexed with human and animal ACE2 proteins.

Specifically for the Omicron variant, both CNN_seq and NN_MM-GBSA models predict improved binding affinity with the *K*_D,app_ ratio values of 1.23 ± 0.58 and 1.25 ± 0.09, respectively. Notably, only 13 out of a total of 25 amino acid change predictions from individual CNN_seq models were improving while the rest were worsening. This alludes to the hypothesis that highly improving towards binding amino acid changes enables neighboring positions to assume binding worsening amino acid changes to evade dominant immune recognition sites.^39,71,87,101–104^ We also calculated the binding free energy Δ*G* of the RBD-hACE2 complex using the Rosetta force-field (see Methods for details). The resulting averaged Δ*G* of the Omicron- and WT-RBD-hACE2 complexes were −36.2 ± 2.4 and −44.3 ± 1.0 kcal/mol, respectively. Experimentally, there does not seem to be a consensus on the relative binding strength between the Omicron variants.^71,87,101^ Surface plasmon resonance analysis by Cameroni et al.^71^ showed that Omicron and Delta bind 2.4- and 1.2-fold stronger than WT, whereas Mannar et al.^101^ suggest 1.5- and 1.4-fold improvement over WT for Omicron and Delta, respectively. Moreover, a combined experimental and simulation study by Wu et al.^87^ concluded that Omicron has a lower binding affinity than Delta and WT. Nevertheless, with as many as 15 amino acid changes in the RBD of the Omicron variant, there could be formation/interruption of multiple residue-residue pairs and substantial structural rearrangement that might be responsible for the inconsistent predictions from different methods.^71,87,101^

As the subvariant of Omicron, BA.2 variant carries one less (G446S), one altered (S371L vs. S371F), and three additional amino acid changes (T376A, D405N, R408S). Notably, these five different mutations were all known to decrease the binding affinity for ACE2.^30^ The CNN_seq model predicts just slightly lower *K*_D,app_ ratio of 1.18 for BA.2 than 1.23 for the original Omicron. In contrast, the B. 1.640.2 variant adds six amino acid changes comprised of four binding-improving (R346S, N394S, E484K, N501Y), one neutral (F490S), and one binding-decreasing (Y449N) mutation,^30^ consistent with the predicted high *K*_D,app_ ratio of 1.73 by the model.

### Using the CNN_seq model built on human data to predict binding with animal ACE2

The inclusion of ACE2 mutations in the training set allows the CNN_seq model to learn how amino acid changes on ACE2 in addition to the RBD could affect the binding affinity change. The model predicted *K*_D,app,WT,animal_/*K*_D,app,WT_ values are summarized in Table 1, where *K*_D,app_ ratio values for cattle, pig, and chicken were smaller than one, indicating weaker binding affinity of SARS-CoV-2 RBD for these animals than human, consistent with the experimental observations.^9,10^ For deer, the WT RBD showed slightly higher binding affinity than human, which seems to agree with the high susceptibility as recently reported.^11–15^ As shown in Table 1, circulating variants generally show enhanced binding affinity for animal ACE2 in all cases in line with corresponding improvements for human. However, the quantitative improvement generally lags behind the one achieved for human ACE2 alluding that adaptation to human ACE2 remains the main driving force of diversity generation. Delta and Kappa variants exhibit slightly higher binding affinity for deer ACE2 than human. Theta, Mu and Alpha+E484K variants show consistently high binding affinity (*K*_D,app_ ratio > 1.3) whereas Gamma, Alpha+E484K+S494P, Iota+L452R, and Iota+L452R+S477N show slightly elevated binding affinity (*K*_D,app_ ratio > 1.0). These CNN_seq model predictions suggest that just as variants achieve tighter binding with hACE2 which may explain gains in infectivity, affinity gains are also predicted against animal ACE2s. The gains in affinity for animal ACE2s appear to track the infection susceptibility of the four assessed animals with deer being very near to human, cattle and horse being significantly less, and chicken only moderately increased from the originally low affinity values.

### Computational scanning for novel variants with up to four amino acid changes using CNN_seq model for human and animals ACE2

On average, it takes approximately 250 ms to predict the *K*_D,app_ ratio for a single variant on an NVIDIA Tesla P100 GPU card. This permits the exhaustive exploration of multiple amino acid changes for possibly further increases in binding affinity with ACE2. Based on the sequence range of the experimentally determined structure of RBD-hACE2 complex,^105^ we focused on RBD variants in the range of residues from 333 to 527. We adopted a hierarchical approach by first exhaustively assessing all single amino acid changes and then selecting the top 20 variants with the highest *K*_D,app_ ratio to exhaustively assess the addition of a second amino acid change. This procedure is repeated until variants with up to four amino acid changes are assessed. This procedure relies on the observation that affinity gains seen so far have been largely, though not exclusively, additive in the contribution of individual amino acid changes in the spike.^106^

Figure 3 compares the CNN_seq predicted and experimental *K*_D,app_ ratio values for variants formed by hACE2 and RBD with single amino acid changes. In total, 195 × 20 = 3,900 RBD variants were scanned, where 3,883 of them have experimental referenced data. Among these variants, 1,344 (34.6%) are in the training set, and the rest 2,539 (65.4%) are unknown to the model. These 2,539 variants are all worsening examples and forms a blind test set, for which the prediction shows a VC% of 94.62% and *r* of 0.90, even higher than the results from training/validation process, further implying the robustness of the model. The binding affinity of circulating variants are reasonably well predicted by the model, showing general agreement with experiments.

**Figure 3.**
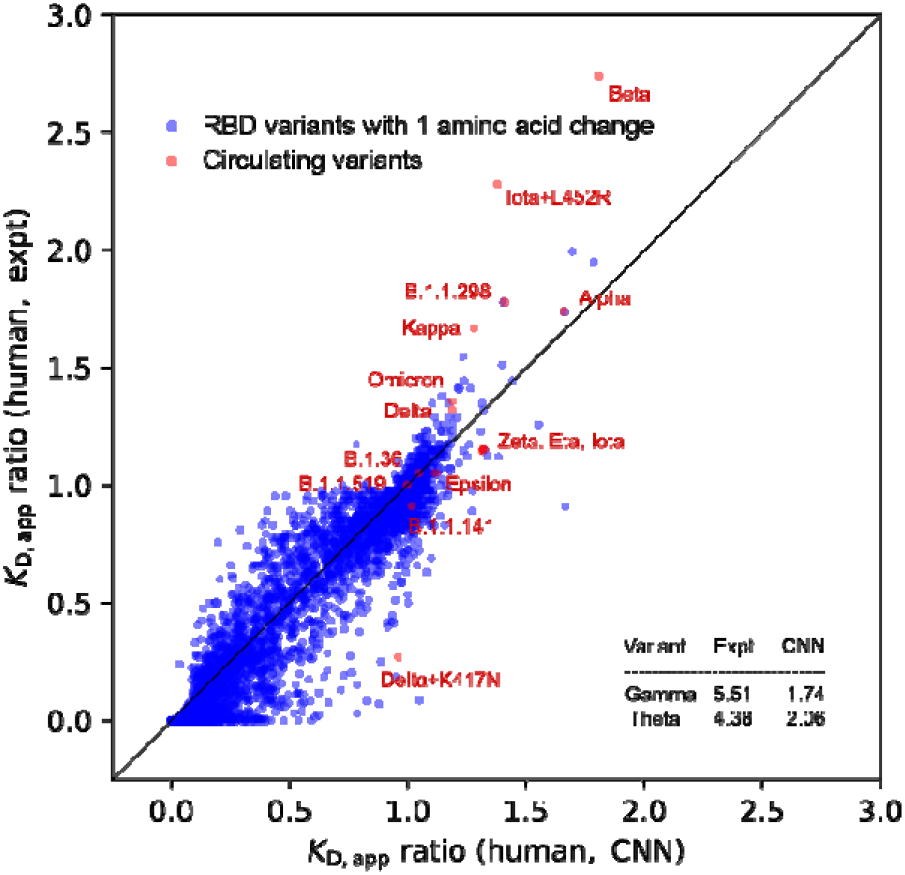
Comparison of *K*_D,app_ ratio between experiments and CNN_seq models predictions from scanning on 3900 variants with one amino acid change. Scanning results are colored in blue, and circulating variants are labeled in red or indicated in the inset table. A diagonal dashed line is drawn to indicate the perfect positive correlation.

To illustrate the variant distribution across the RBD sites under investigation, in Figure 4 we constructed the binding affinity change heatmaps for all 3,900 variants with one amino acid changes followed the pictorial style introduced in Starr et al.^30^ Each one of the stripes represents the complete scanning results for one of the species, where the horizontal and vertical axes indicate the RBD sites and amino acid, respectively. The small squares are colored according to the CNN_seq predicted *K*_D,app_ ratio values, with red for lower affinity, blue for higher affinity, and white for neutral or similar affinity. The general patterns of the heatmap for the experimental data (Figure 4A) are well captured in the CNN_seq predictions (Figure 4B) for the human case, in line with the VC% and *r* values obtained from data shown in Figure 3A.

**Figure 4.**
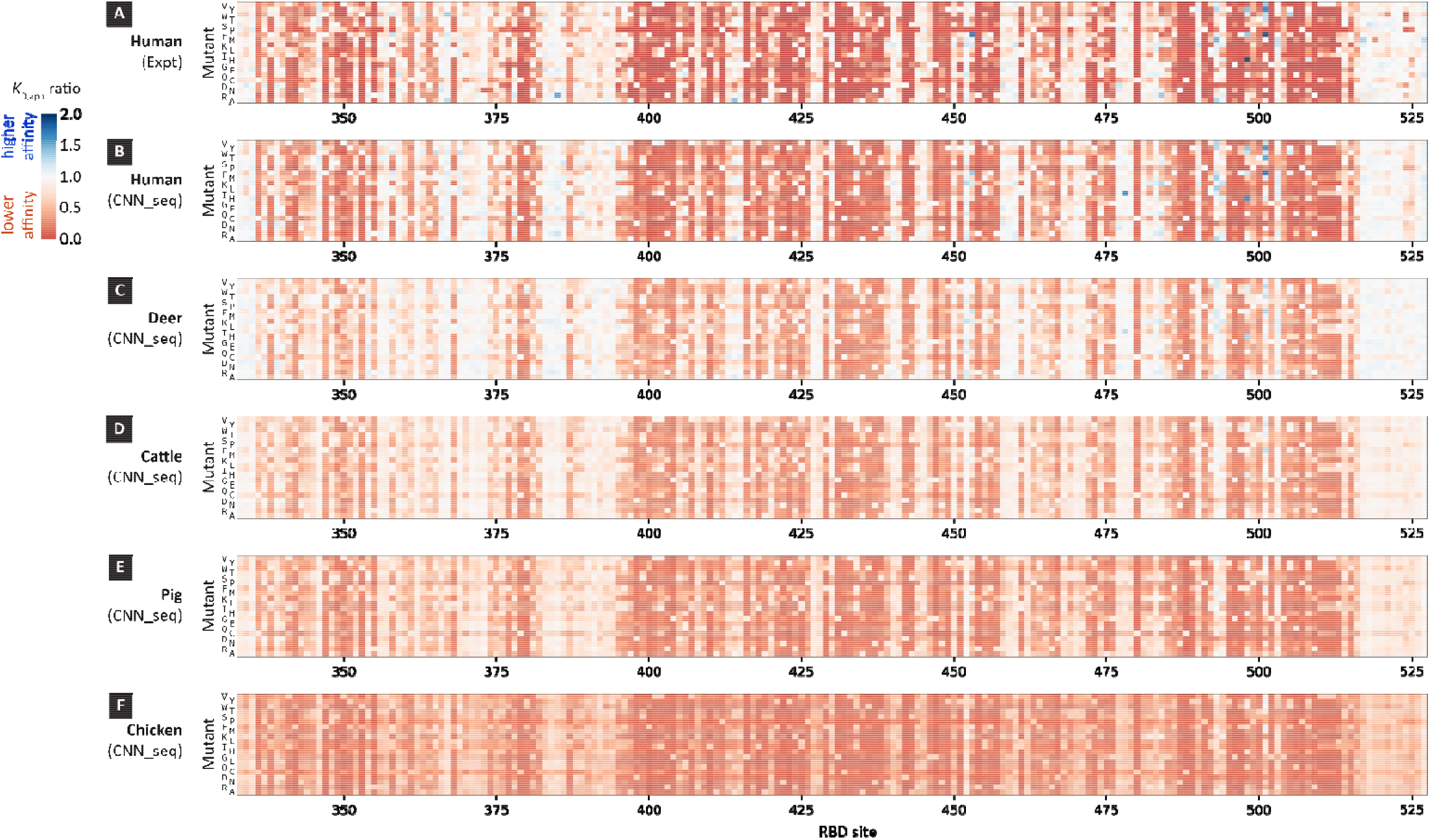
Binding affinity change heatmaps of RBD variants for ACE2 proteins from human and animal hosts. Experimental values and CNN_seq predictions for human are shown in (A) and (B), and CNN_seq predictions for animals are depicted in (C) deer, (D) cattle, (E) pig, (F) chicken. Squares are colored according to the *K*_D,app_ ratio values.

To examine the effects of RBD mutations on the change of binding affinity for animal ACE2 proteins, we further scanned the 3,900 RBD variants with one amino acid changes for all four animal hosts. As illustrated in Figure 4C to Figure 4F, the heat maps largely resemble the ones for human in proportion to the sequence similarity between their ACE2 proteins. In Figure 5 we compare the deer vs. human *K*_D,app_ ratio values for all scanned 3,900 data and circulating variants. Results for other animals are shown in Figure S2 in *SI Appendix*. Correlations are observed between human and all animals RBD variants (see Figure 4).

**Figure 5.**
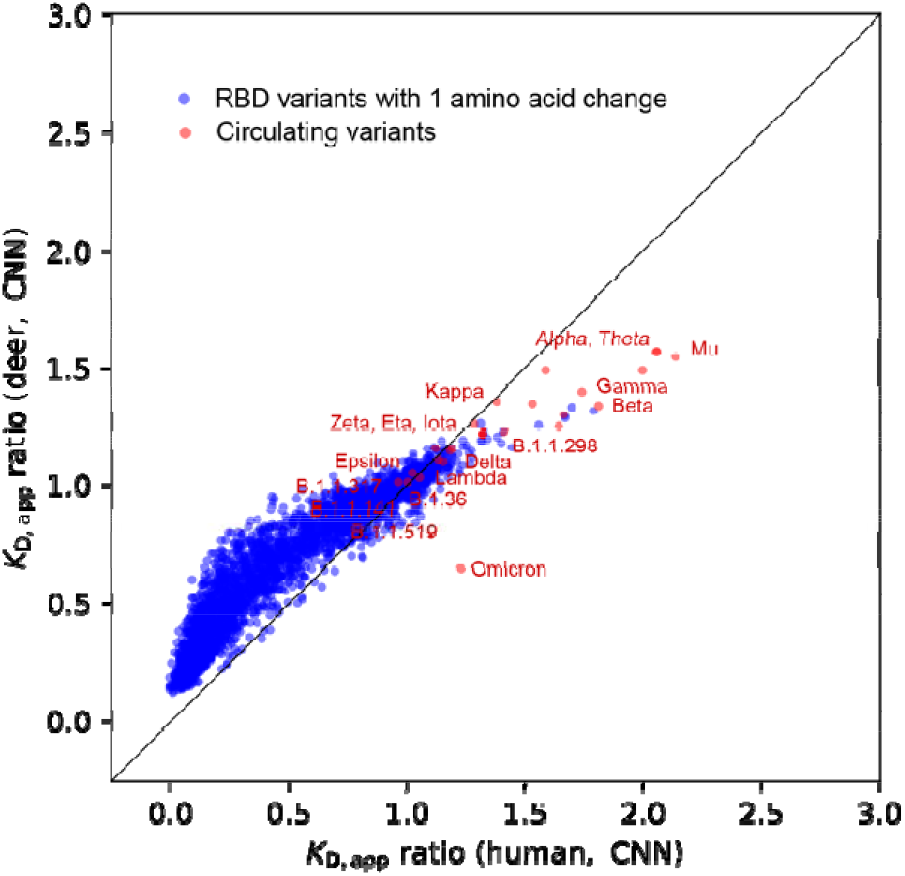
Comparison of binding affinities of RBD variants for ACE2 proteins of human and deer. All the RBD variants with single amino acid changes are colored in blue, circulating variants are labeled in red. A diagonal line is drawn to indicate the perfect positive correlation.

With the scanning results, we could examine whether adaptation of the virus for increased binding with animals would cause concomitant increases to binding with human ACE2 (animal/human converging amino acid changes) or decreased fitness. The top deer/human converging and diverging amino acid changes are tabulated in Table S2 in *SI Appendix*. Among the deer/human converging amino acid changes, there are many candidates that show significant binding affinity enhancement for both deer and human including N501Y, E484K, Y453F, L452R that are contained in multiple circulating variants. For deer/human diverging amino acid changes, most candidates show moderated binding affinity increase for deer and mild decrease for human. Notably, change A522S was identified in B.1.1.317 variant, which circulated in Russia, UK, and Thailand in late 2020.^107^

The effect of multiple amino acid changes (see Table 2) is difficult to interpret as they may cause significant re-organization of the RBD. It is unclear whether continually improving binding affinity would translate to increased infectivity as other biological processes underpinning productive cell infection and proliferation may become limiting (e.g., furin cleavage, RBD presentation, or internalization efficiency). However, the computationally predicted potential to reach even more increased binding affinity may suggest a still untapped potential of SARS-CoV-2 to undergo additional immune evasion changes whose potentially detrimental effect on binding could be ameliorated by a set of improving amino acid changes. As seen in Table 2, the effect of adding amino acids changes is not always additive. There are many instances where the best double change variant combines an amino acid change with very high affinity and another with mild or even lower affinity. Tables S3, S4, S5 and S6 list corresponding results for different animals in SI Appendix.

**Table 2.**
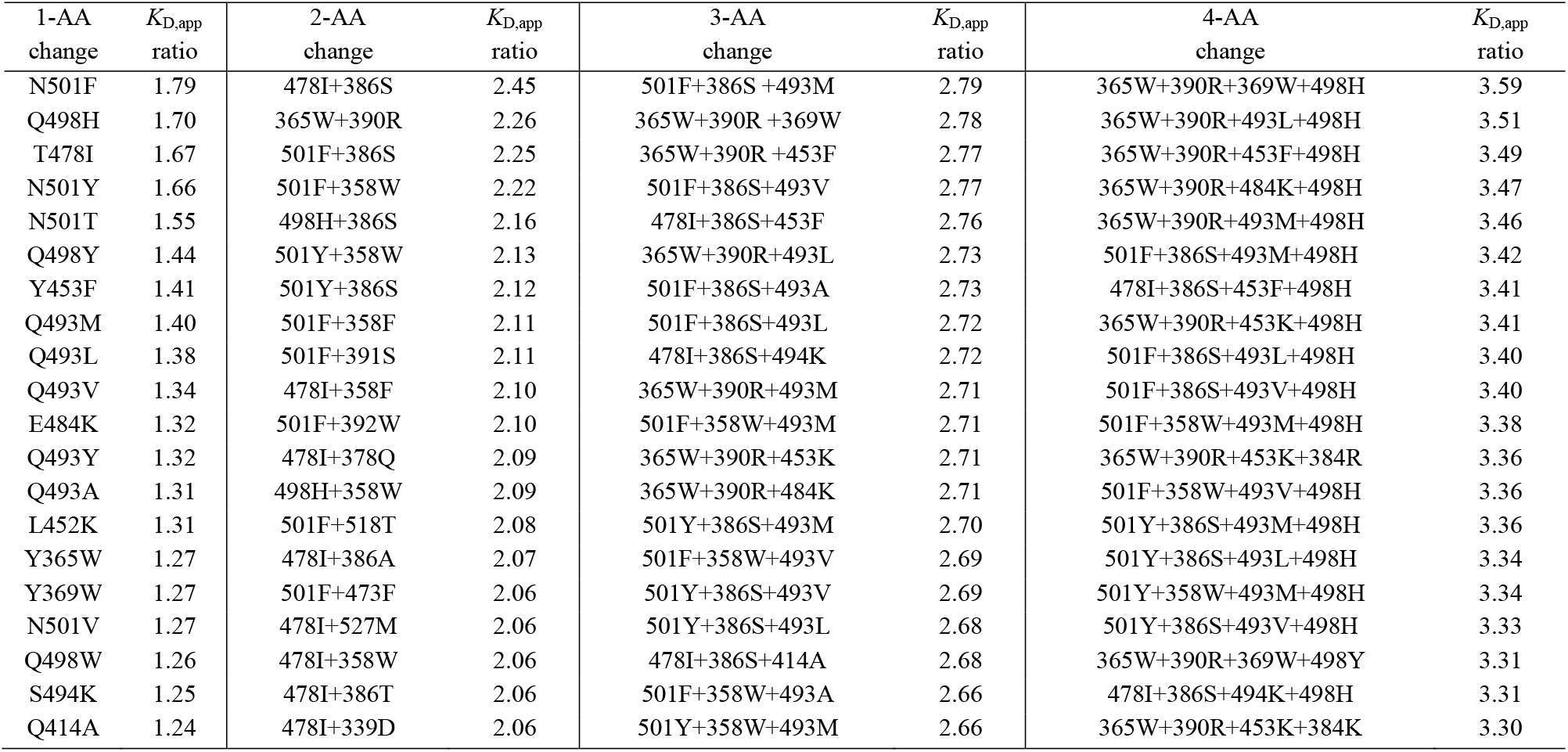
Scanning results for human with up to 4 amino acid changes on SARS-CoV-2 RBD.

| 1-AA<br>change | $K_{D,app}$<br>ratio | 2-AA<br>change | $K_{D,app}$<br>ratio | 3-AA<br>change | $K_{D,app}$<br>ratio | 4-AA<br>change | $K_{D,app}$<br>ratio |
| --- | --- | --- | --- | --- | --- | --- | --- |
| N501F | 1.79 | 478I+386S | 2.45 | 501F+386S+493M | 2.79 | 365W+390R+369W+498H | 3.59 |
| Q498H | 1.70 | 365W+390R | 2.26 | 365W+390R+369W | 2.78 | 365W+390R+493L+498H | 3.51 |
| T478I | 1.67 | 501F+386S | 2.25 | 365W+390R+453F | 2.77 | 365W+390R+453F+498H | 3.49 |
| N501Y | 1.66 | 501F+358W | 2.22 | 501F+386S+493V | 2.77 | 365W+390R+484K+498H | 3.47 |
| N501T | 1.55 | 498H+386S | 2.16 | 478I+386S+453F | 2.76 | 365W+390R+493M+498H | 3.46 |
| Q498Y | 1.44 | 501Y+358W | 2.13 | 365W+390R+493L | 2.73 | 501F+386S+493M+498H | 3.42 |
| Y453F | 1.41 | 501Y+386S | 2.12 | 501F+386S+493A | 2.73 | 478I+386S+453F+498H | 3.41 |
| Q493M | 1.40 | 501F+358F | 2.11 | 501F+386S+493L | 2.72 | 365W+390R+453K+498H | 3.41 |
| Q493L | 1.38 | 501F+391S | 2.11 | 478I+386S+494K | 2.72 | 501F+386S+493L+498H | 3.40 |
| Q493V | 1.34 | 478I+358F | 2.10 | 365W+390R+493M | 2.71 | 501F+386S+493V+498H | 3.40 |
| E484K | 1.32 | 501F+392W | 2.10 | 501F+358W+493M | 2.71 | 501F+358W+493M+498H | 3.38 |
| Q493Y | 1.32 | 478I+378Q | 2.09 | 365W+390R+453K | 2.71 | 365W+390R+453K+384R | 3.36 |
| Q493A | 1.31 | 498H+358W | 2.09 | 365W+390R+484K | 2.71 | 501F+358W+493V+498H | 3.36 |
| L452K | 1.31 | 501F+518T | 2.08 | 501Y+386S+493M | 2.70 | 501Y+386S+493M+498H | 3.36 |
| Y365W | 1.27 | 478I+386A | 2.07 | 501F+358W+493V | 2.69 | 501Y+386S+493L+498H | 3.34 |
| Y369W | 1.27 | 501F+473F | 2.06 | 501Y+386S+493V | 2.69 | 501Y+358W+493M+498H | 3.34 |
| N501V | 1.27 | 478I+527M | 2.06 | 501Y+386S+493L | 2.68 | 501Y+386S+493V+498H | 3.33 |
| Q498W | 1.26 | 478I+358W | 2.06 | 478I+386S+414A | 2.68 | 365W+390R+369W+498Y | 3.31 |
| S494K | 1.25 | 478I+386T | 2.06 | 501F+358W+493A | 2.66 | 478I+386S+494K+498H | 3.31 |
| Q414A | 1.24 | 478I+339D | 2.06 | 501Y+358W+493M | 2.66 | 365W+390R+453K+384K | 3.30 |

## 3. Discussion

The introduced CNN_seq model overcomes the computational barriers associated with our previous NN_MD-MMGBSA procedure^58^ and at the same time unlocks the opportunity to consider amino acid changes throughout the entire RBD as well as ACE2 receptor during model training. The enlarged (by about 80-fold) training set affords the CNN_seq model to learn more effectively and potentially capture distal correlations. Meanwhile, the improved efficiency and accessibility of feature encoding expand the range of possible variants that can be assessed for binding affinity changes with human and animal ACE2s.

Focusing on the Omicron variant as an exemplar application, the CNN_seq model delineated the role of mutations as binding improving and/or immune escaping^108,109^. In addition, given that it is intractable to exhaustively assess the binding affinity changes in response to two or more amino acid changes experimentally, the CNN_seq model could serve as a computational alternative for the *a priori* assessment of possible epistasis and synergistic effects.^104^ In addition, pre-calculation of very high binding affinity variants involving multiple amino acid changes using CNN and continuous surveillance of circulating variant databases could offer an alert system whenever an emerging variant is becoming sequence-adjacent to a computationally predicted problematic variant. This could become part of a surveillance system to flag variant amino acid changes seen in human or animal hosts that upon one or more additional changes could become a particularly problematic variant.

## 4. Methods

### Homology modeling using Swiss-Model

Homology modelling was performed using SWISS-Model. For each animal, the sequence of specific ACE2 was concatenated with that of SARS-CoV-2 spike RBD, and loaded into the server interface. A representative subset of the template structures can be automatically extracted from the database through multiway alignment.

### Refinement of homology models using molecular dynamics simulations

The structures of homology models were refined through MD simulations. Each RBD-ACE2 complex structure was first prepared using protein preparation wizard of Maestro in Schrödinger suite (v2019.4), and subsequently solvated in an orthorhombic box with 10 Å buffer in all three dimensions. The TIP3P model^110^ was chosen for water molecules and the whole system was neutralized by adding Na^+^ and Cl^−^ ions to reach a salt concentration of 0.15 M. The Amber99SB-ILDN force field^111^ was used to account for the interactions of proteins, while the interactions between protein and water molecules were automatically generated by the Desmond force field building tool Viparr. The solvated system was minimized and equilibrated following the default relaxation protocol of Desmond^112^, and a 100-ns production run was subsequently conducted to fully relax and optimize the structure of the RBD-ACE2 complex. The production run was performed in isothermal-isobaric (NPT) ensemble with periodic boundary conditions applied in all three dimensions, and a temperature of 300 K and a pressure of 1.0 atm was maintained. A time step of 2.0 fs was set to integrate the equations of motion, particle mesh Ewald method was used to describe the long-range interactions, and a cutoff distance of 9.0 Å was applied in the calculation of non-bonded short-range interactions. The convergence of the system was verified by monitoring the RMSD of the simulation run. For all RBD-ACE2 complexes, the RMSD values kept below 1 Å in the final 100-ns of the production run, suggesting convergence of the structure optimization.

### Generate distance map based on structure using Protein Contact Maps

Since the base structure of each RBD-ACE2 complex requires only one optimization, we prepared the protein contact map and used it as the constant input during CNN model construction. The protein contact map can be generated by utilizing the Protein Contact Maps tool developed by Benjamin et al. Each optimized RBD-ACE2 complex structure was loaded into the online server and the outputs were the contact map and the distance map associated with the individual input structure. The obtained binary two-dimensional matrix distance map is indexed following the provided protein sequence and will be accessed in CNN feature encoding process.

### Identification of protein-protein interfacial contacts using PDBePISA

Protein Data Bank in Europe (PDBe) offers a web tool PISA (i.e., Proteins, Interfaces, Structures and Assemblies) to help access the interfacial interactions between proteins. By loading the optimized RBD-ACE2 complex structure into the PISA server, the interfacial contact information can be promptly generated. Among the rich output content, we take the interfacial residues, the contact pairs and the distances for hydrogen bonds or salt bridges. This information is used in CNN feature encoding process to label whether a specific residue is at interface or involved in hydrogen bonds and salt bridges, such that the weight of the feature could be adjusted independently.

### Dataset generation

The total number of all available ACE2 and RBD variants sum up to 93,669. There are predominantly more worsening (97.02%) than improving (2.38%) or neutral (0.60%) variants. Absorbing all of the variants will create imbalanced dataset that may result in models with poor performance. To tackle this problem, we chose to include all the improving variants with up to six amino acid changes from the RBD and ACE2 variants, and randomly picking three times as many worsening or neutral variants accordingly. This process creates a dataset of 8,440 variants where the number (percentage) of improving, worsening, neutral variants are 2,212 (26.21%), 5,667 (67.14%), 561 (6.65%), respectively. This dataset contains 6,105 (72.10%) variants with RBD mutations, 2335 (27.90%) variants with ACE2 mutations, 1341 (15.89%) variants with single amino acid changes, and 7099 (84.11%) variants with multiple amino acid changes.

### Hyperparameter optimization

The Bayesian hyperparameter optimization was performed by utilizing the Hyperopt package (http://hyperopt.github.io/hyperopt) to achieve optimal performance of the CNN_seq model. The loss function for minimization was defined as:

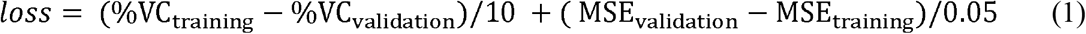

The loss function numerically evaluates the gap in performance of CNN_seq model on training and validation set, and the constants 10 and 0.05 were chosen to match the difference in %VC and MSE with a roughly equal contribution. A total of 50 iterations of optimization were performed to achieve the final set of hyperparameters that are summarized in Table S7 in *SI Appendix*.

### Model training, evaluation, and prediction

The model was trained with the objective function as the MSE between predicted and target *K*_D,app_ ratio values. The Adam optimizer^113^ was used to perform the backpropagation and the training was performed for 5,000 epochs including the entire training data in each batch. To measure the strength of correlation between experimental and predicted *K*_D,app_ ratios, we calculate the Pearson correlation coefficient *r*, which is defined as:

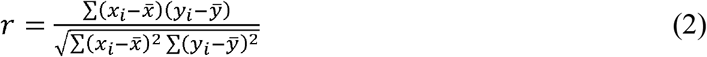

where *x_i_* and *y_i_* are the target and predicted value for the *i*^th^ sample, and 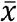 and 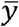 are the mean value for all *x_i_* targets and *y_i_* predictions. Note that, due to classification ambiguity, variants with either target value or predicted value of *K*_D,app_ ratio equals to 1 were excluded during the calculation of %VC, but all variants were included during the calculation of *r*.

The final predictions of CNN_seq model in Table 1 are calculated using a single model trained on 100% of the variants selected from the full database. The variants in Table 1 were excluded from the training data so that a fair evaluation can be made.

### Rosetta calculations for ΔΔ*G*_bind_ prediction

To computationally assess the binding affinity of the Omicron-RBD with hACE2, we used rigorous molecular-mechanics based calculations using the Rosetta force-field. Assuming that the Omicron-RBD still binds to hACE2 at the same binding site as the WT-RBD, we first made the initial complex of Omicron-RBD-hACE2 by making all the 15 amino-acid changes.^6^ To account for structural re-arrangements of the mutated RBD in the complex, the initial complex was subject to 100 independent *Relax* trajectories. Harmonic constraints were used to prevent the structure from deviating significantly from the crystal structure. At the end of *Relax*, a gradient minimization is performed using *lbfgs_armijo* algorithm for 2000 steps after which the relevant metrics of binding were calculated using *InterfaceAnalyzer*. The binding energy, Δ*G* of the Omicron-RBD-hACE2 complex was then calculated as the average of *dG_separated* scores obtained from the 100 *Relax* simulations.

## Data Availability

Computational codes were developed in Python using the PyTorch library. All data pertaining to the results discussed in the paper are available either in the main text and *SI Appendix*. Relevant simulation codes for generating the models are deposited in the GitHub repository (https://github.com/maranasgroup/CNN_seq_CoV2).

## Acknowledgements

This activity was primarily supported by the United States Department of Agriculture (USDA) NIFA Award 2020-67015-32175 and also partially enabled by funding provided by The Center for Bioenergy Innovation a U.S. Department of Energy Research Center supported by the Office of Biological and Environmental Research in the DOE Office of Science (DE-AC05-000R22725).

Computations for this research were performed on the Pennsylvania State University’s Institute for Computational and Data Sciences’ Roar supercomputer. We also acknowledge Seed grant funding from the Penn State Huck Institutes of life sciences (to SVK).

Kurt Vandegrift was partially supported by a National Science Foundation Ecology and Evolution of Infectious Diseases program grant (# 1619072) as well as National Institute of Allergy and Infectious Diseases, National Institutes of Health grants (#1R21AI156406-01 & #1R01AI134911).

## Author Approvals

All the authors approve of the work.

## Appendix A. Supplementary data

Supplementary data to this article are given in “Supplementary Information.pdf”

