## Supplementary Information for "A CNN model for predicting binding affinity changes between SARS-CoV-2 spike RBD variants and ACE2 homologues"

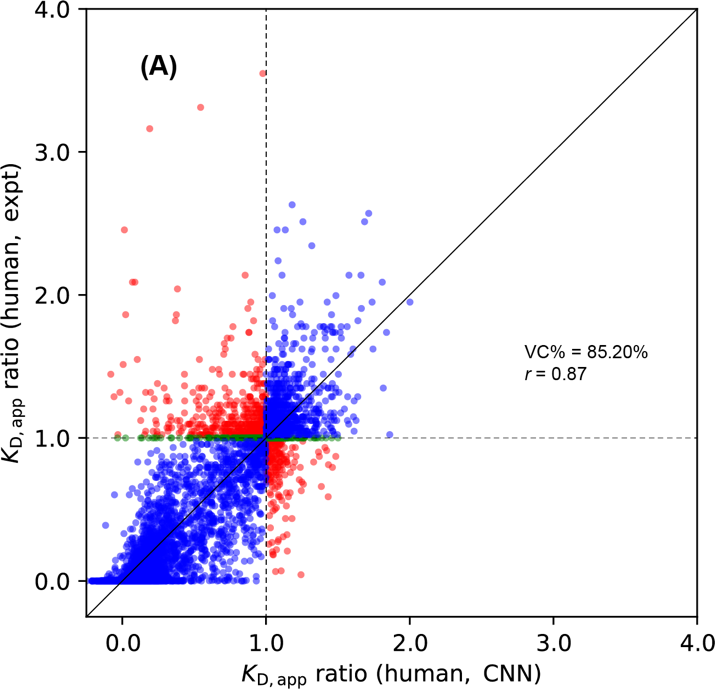

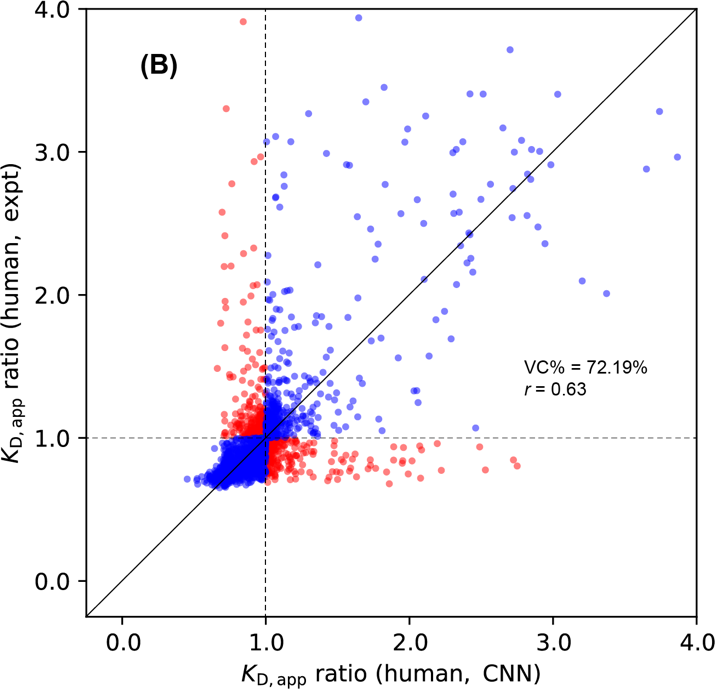

**Figure S1.** Comparison of *K*_D,app_ ratio between experiments and CNN_seq models predictions from five-fold cross-validation tests on variants with amino acid changes on (A) RBD and (B) ACE2 variants. Correctly classified variants are colored in blue, incorrectly classified variants are colored in red, and variants with unchanged binding affinities are colored in green. Horizontal and vertical dashed lines are drawn to indicate the dividing line where *K*_D,app_ ratio equals to 1, and a diagonal solid line is drawn to indicate perfect correlation.

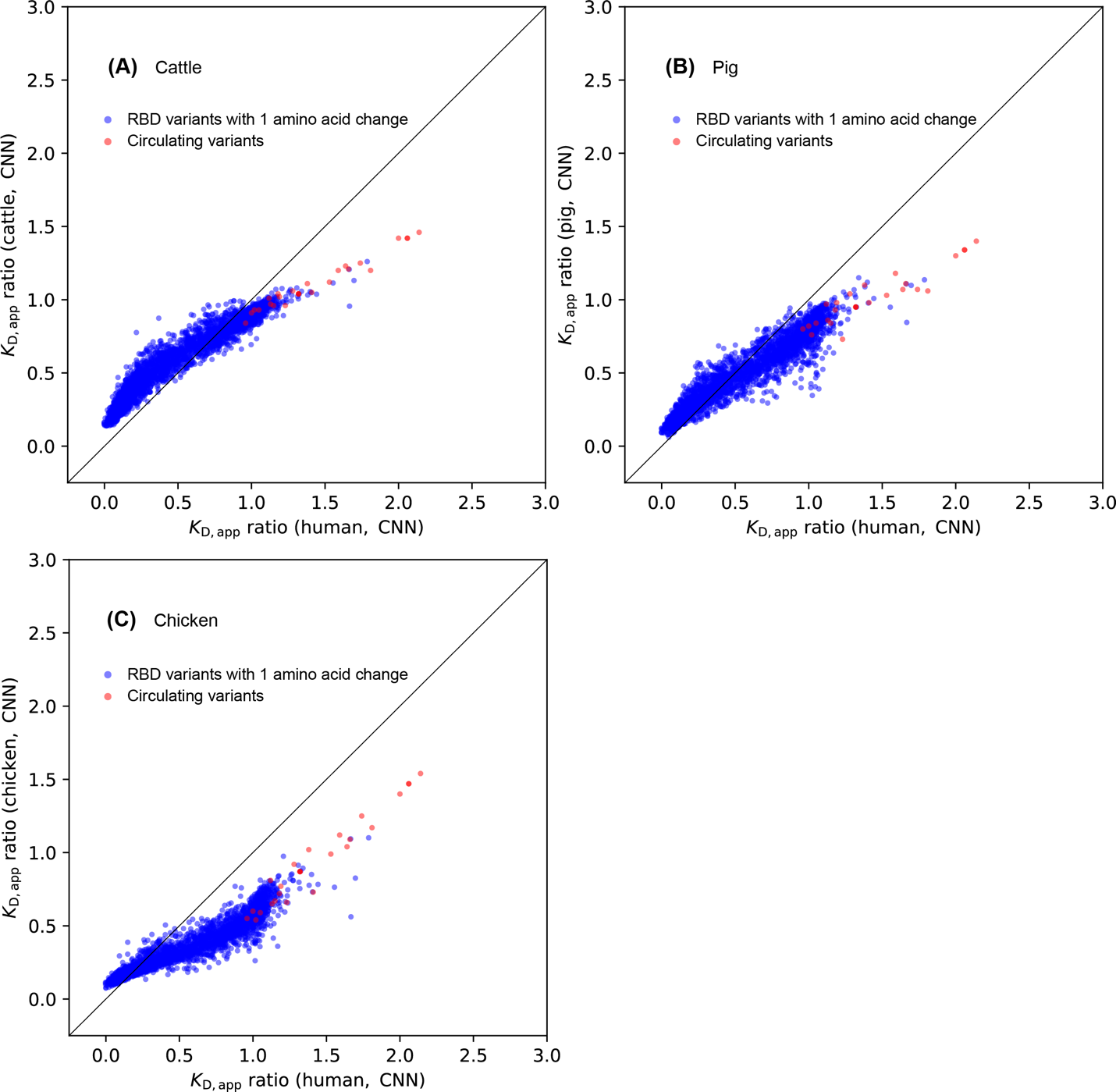

**Figure S2.** Comparison of binding affinities of RBD variants for ACE2 proteins of human and (A) cattle, (B) pig, and (C) chicken. All the RBD variants with single amino acid changes are colored in blue, circulating variants are labeled in red. A diagonal line is drawn to indicate the perfect positive correlation.

**Table S1.** Comparison of *K*_D,app_ ratio from experiments, CNN_seq, and NN_MM-GBSA model predictions on variants with multiple AA changes.

| **Amino acid change(s) in RBD** |  | ***K*_D,app_ ratio** |  |
| --- | --- | --- | --- |
|  | **Experiment** | **CNN_seq** | **NN_MMGBSA** |
| E484K+N501Y | 4.38 | 2.06 | 1.22 |
| L452R+T478K | 1.32 | 1.19 | 1.12 |
| L452R+E484K | 2.28 | 1.38 | 1.22 |
| L452R+E484Q | 1.67 | 1.28 | 1.21 |
| K417T+E484K | 0.49 | 0.85 | 1.25 |
| K417N+L452R | 0.61 | 0.97 | 0.83 |
| K417N+E484K | 0.28 | 0.61 | 1.25 |
| K417N+N501Y | 1.01 | 1.42 | 1.25 |
| L452R+N501Y | 7.65 | 1.47 | 1.25 |
| Y453F+N439K | 1.04 | 1.36 | 1.25 |
| K417V+N439K | 1.00 | 0.54 | 1.20 |
| H374N+H378N | 0.21 | 0.41 | 1.11 |
| K417N+E484K+N501Y | 2.74 | 1.81 | 1.22 |
| K417T+E484K+N501Y | 5.51 | 1.74 | 1.22 |
| K417N+L452R+E484K+N501Y | 2.33 | 1.71 | 1.25 |

**Table S2.** Top 20 deer/human converging and diverging amino acid changes.

| Converging amino acid change | *K*_D,app_ ratio (deer) | *K*_D,app_ ratio (human) | Diverging  amino acid change | *K*_D,app_ ratio (deer) | *K*_D,app_ ratio (human) |
| --- | --- | --- | --- | --- | --- |
| 498H | 1.33 | 1.70 | 354D | 1.10 | 0.98 |
| 501F | 1.32 | 1.79 | 391S | 1.09 | 0.87 |
| 501Y | 1.30 | 1.66 | 354A | 1.08 | 0.91 |
| 478I | 1.29 | 1.67 | 391Q | 1.06 | 0.94 |
| 452K | 1.26 | 1.31 | 372N | 1.06 | 0.99 |
| 501T | 1.25 | 1.55 | 383R | 1.06 | 0.99 |
| 453F | 1.23 | 1.41 | 391N | 1.06 | 0.90 |
| 484K | 1.22 | 1.32 | 351F | 1.06 | 1.00 |
| 493L | 1.21 | 1.38 | 503Q | 1.06 | 0.99 |
| 493V | 1.20 | 1.34 | 372K | 1.06 | 0.95 |
| 493F | 1.19 | 1.24 | 371D | 1.05 | 1.00 |
| 493A | 1.19 | 1.31 | 384N | 1.05 | 0.99 |
| 503R | 1.19 | 1.15 | 391T | 1.05 | 0.84 |
| 493Y | 1.19 | 1.32 | 371Q | 1.05 | 0.94 |
| 493M | 1.18 | 1.40 | 354H | 1.05 | 0.93 |
| 494K | 1.17 | 1.25 | 455V | 1.05 | 0.95 |
| 414A | 1.17 | 1.24 | 478N | 1.05 | 0.99 |
| 354S | 1.17 | 1.11 | 526K | 1.04 | 0.99 |
| 498Y | 1.16 | 1.44 | 337E | 1.04 | 0.99 |
| 452A | 1.16 | 1.16 | 501A | 1.04 | 0.92 |

**Table S3.** Scanning results for deer with up to 4 AA changes on SARS-CoV-2 RBD.

| 1-AA change | *K*_D,app_ ratio deer | *K*_D,app_ ratio human | 2-AA  change | *K*_D,app_ ratio deer | *K*_D,app_ ratio human | 3-AA  change | *K*_D,app_ ratio deer | *K*_D,app_ ratio human | 4-AA  change | *K*_D,app_ ratio deer | *K*_D,app_ ratio human |
| --- | --- | --- | --- | --- | --- | --- | --- | --- | --- | --- | --- |
| Q498H | 1.33 | 1.70 | 501F+391S | 1.61 | 2.11 | 498H+369W+390R | 2.08 | 2.73 | 498H+365W+384R+390R | 2.64 | 3.45 |
| N501F | 1.32 | 1.79 | 498H+518T | 1.60 | 2.05 | 498H+369W+390W | 2.00 | 2.58 | 498H+365W+369W+390R | 2.62 | 3.59 |
| N501Y | 1.30 | 1.66 | 498H+391S | 1.57 | 1.83 | 478I+501F+391S | 1.99 | 2.88 | 498H+369W+380W+390R | 2.51 | 2.89 |
| T478I | 1.29 | 1.67 | 452K+339N | 1.57 | 1.62 | 478I+498H+518T | 1.98 | 2.89 | 498H+365W+384R+390W | 2.45 | 3.20 |
| L452K | 1.26 | 1.31 | 501Y+391S | 1.57 | 1.93 | 478I+501Y+391S | 1.97 | 2.70 | 498H+369W+382W+390R | 2.44 | 2.97 |
| N501T | 1.25 | 1.55 | 498H+518G | 1.56 | 1.89 | 498H+384R+390W | 1.96 | 2.49 | 498H+527R+369W+390R | 2.42 | 3.16 |
| Y453F | 1.23 | 1.41 | 498H+358W | 1.55 | 2.09 | 478I+498H+391S | 1.96 | 2.62 | 498H+362W+369W+390R | 2.41 | 2.93 |
| E484K | 1.22 | 1.32 | 501F+518G | 1.55 | 2.03 | 478I+498H+518G | 1.95 | 2.72 | 498H+527W+369W+390R | 2.41 | 3.22 |
| Q493L | 1.21 | 1.38 | 501F+391T | 1.54 | 2.00 | 498H+384R+390R | 1.95 | 2.44 | 493L+498H+369W+390R | 2.39 | 3.15 |
| Q493V | 1.20 | 1.34 | 501Y+358W | 1.54 | 2.13 | 478I+500Y+501T | 1.95 | 2.71 | 498H+365W+369W+390W | 2.38 | 3.19 |
| Q493F | 1.19 | 1.24 | 500Y+501T | 1.54 | 1.90 | 478I+498H+358W | 1.94 | 2.94 | 498H+527K+369W+390R | 2.38 | 3.13 |
| Q493A | 1.19 | 1.31 | 501F+358W | 1.54 | 2.22 | 478I+498H+358F | 1.93 | 2.91 | 498H+380W+384R+390R | 2.34 | 2.63 |
| V503R | 1.19 | 1.15 | 498H+518S | 1.54 | 1.95 | 478I+498H+369W | 1.93 | 2.92 | 498H+367Q+369W+390W | 2.34 | 3.09 |
| Q493Y | 1.19 | 1.32 | 498H+369W | 1.53 | 2.04 | 478I+501Y+358W | 1.92 | 2.96 | 498H+365F+369W+390R | 2.33 | 3.12 |
| Q493M | 1.18 | 1.40 | 498H+384R | 1.53 | 1.89 | 478I+501F+391T | 1.92 | 2.77 | 498H+524W+369W+390R | 2.33 | 2.89 |
| S494K | 1.17 | 1.25 | 501F+391R | 1.53 | 1.99 | 478I+498H+518S | 1.92 | 2.78 | 498H+365F+384R+390R | 2.33 | 3.03 |
| Q414A | 1.17 | 1.24 | 498H+358F | 1.53 | 2.03 | 478I+501F+518G | 1.92 | 2.84 | 498H+362F+369W+390R | 2.33 | 2.85 |
| N354S | 1.17 | 1.11 | 498H+392W | 1.53 | 2.02 | 453F+501Y+391S | 1.91 | 2.45 | 493L+498H+384R+390W | 2.32 | 2.98 |
| Q498Y | 1.16 | 1.44 | 452K+339D | 1.53 | 1.65 | 478I+498H+392W | 1.91 | 2.87 | 498H+367Q+384R+390W | 2.32 | 2.95 |
| L452A | 1.16 | 1.16 | 498H+391R | 1.52 | 1.81 | 484K+501F+391S | 1.91 | 2.47 | 493L+498H+369W+390W | 2.32 | 3.02 |

**Table S4.** Scanning results for cattle with up to 4 AA changes on SARS-CoV-2 RBD.

| 1-AA change | *K*_D,app_ ratio cattle | *K*_D,app_ ratio human | 2-AA  change | *K*_D,app_ ratio cattle | *K*_D,app_ ratio human | 3-AA  change | *K*_D,app_ ratio cattle | *K*_D,app_ ratio human | 4-AA  change | *K*_D,app_ ratio cattle | *K*_D,app_ ratio human |
| --- | --- | --- | --- | --- | --- | --- | --- | --- | --- | --- | --- |
| N501F | 1.26 | 1.79 | 478I+501F | 1.50 | 2.58 | 498H+365W+390R | 1.91 | 3.11 | 498H+365W+384R+390R | 2.30 | 3.45 |
| N501Y | 1.21 | 1.66 | 501F+391S | 1.46 | 2.11 | 498H+501F+391S | 1.87 | 2.76 | 498H+501F+382W+390R | 2.28 | 3.21 |
| Q498H | 1.13 | 1.70 | 365W+390R | 1.45 | 2.26 | 448G+498H+501Y | 1.85 | 2.77 | 498H+365W+384K+390R | 2.27 | 3.41 |
| N501T | 1.11 | 1.55 | 478I+501Y | 1.43 | 2.46 | 440G+498H+501Y | 1.85 | 2.68 | 453F+498H+365W+390R | 2.26 | 3.49 |
| L452K | 1.08 | 1.31 | 501F+341G | 1.43 | 1.79 | 478I+498H+501F | 1.84 | 3.16 | 498H+501F+384R+390R | 2.25 | 3.19 |
| N501W | 1.08 | 1.21 | 501F+367Q | 1.42 | 2.02 | 501F+527W+390R | 1.84 | 2.63 | 453K+498H+365W+390R | 2.25 | 3.41 |
| N501M | 1.07 | 1.17 | 501F+518G | 1.42 | 2.03 | 498H+501F+391T | 1.83 | 2.67 | 501F+527W+384R+390R | 2.25 | 3.11 |
| N501V | 1.07 | 1.27 | 501F+358W | 1.42 | 2.22 | 478I+498T+501Y | 1.82 | 3.10 | 448G+493M+498H+501Y | 2.25 | 3.37 |
| Q498W | 1.06 | 1.26 | 473F+501F | 1.42 | 2.06 | 498H+501F+518G | 1.82 | 2.69 | 498H+365W+369W+390R | 2.24 | 3.59 |
| Q493V | 1.06 | 1.34 | 501F+390R | 1.42 | 2.02 | 498H+501F+390R | 1.82 | 2.67 | 493M+498H+501F+391S | 2.22 | 3.31 |
| Q493M | 1.06 | 1.40 | 501F+339D | 1.41 | 2.05 | 498H+501F+358W | 1.81 | 2.84 | 493L+498H+501F+391S | 2.22 | 3.27 |
| Y453F | 1.05 | 1.41 | 501F+391R | 1.41 | 1.99 | 498H+501F+518T | 1.81 | 2.76 | 448G+493L+498H+501Y | 2.22 | 3.31 |
| S494K | 1.05 | 1.25 | 501F+391T | 1.41 | 2.00 | 498H+501F+391R | 1.79 | 2.63 | 493L+498H+365W+390R | 2.22 | 3.51 |
| N501H | 1.04 | 1.14 | 440G+501Y | 1.41 | 1.96 | 478I+498H+501Y | 1.79 | 3.09 | 498H+365W+380W+390R | 2.22 | 3.24 |
| E484K | 1.04 | 1.32 | 448G+501Y | 1.40 | 2.03 | 450G+498H+501F | 1.79 | 2.65 | 498H+339D+365W+390R | 2.21 | 3.39 |
| Q498Y | 1.04 | 1.44 | 501F+392W | 1.40 | 2.10 | 498T+501F+391S | 1.79 | 2.60 | 484K+498H+365W+390R | 2.21 | 3.47 |
| Q493L | 1.03 | 1.38 | 501F+518T | 1.40 | 2.08 | 478I+498A+501Y | 1.78 | 3.04 | 448G+493V+498H+501Y | 2.21 | 3.33 |
| Y365W | 1.03 | 1.27 | 450G+501F | 1.39 | 1.97 | 498H+501F+392W | 1.78 | 2.72 | 501F+527W+384K+390R | 2.21 | 3.09 |
| Q493A | 1.02 | 1.31 | 501F+367N | 1.39 | 2.00 | 498H+501F+367Q | 1.78 | 2.63 | 440G+493M+498H+501Y | 2.21 | 3.24 |
| L452Q | 1.02 | 1.17 | 501F+358F | 1.39 | 2.11 | 473F+498H+501F | 1.78 | 2.67 | 498H+501F+384W+390R | 2.20 | 3.15 |

**Table S5.** Scanning results for pig with up to 4 AA changes on SARS-CoV-2 RBD.

| 1-AA change | *K*_D,app_ ratio pig | *K*_D,app_ ratio human | 2-AA  change | *K*_D,app_ ratio pig | *K*_D,app_ ratio human | 3-AA  change | *K*_D,app_ ratio pig | *K*_D,app_ ratio human | 4-AA  change | *K*_D,app_ ratio pig | *K*_D,app_ ratio human |
| --- | --- | --- | --- | --- | --- | --- | --- | --- | --- | --- | --- |
| Q493V | 1.15 | 1.34 | 501F+391S | 1.48 | 2.11 | 494K+501F+391S | 1.82 | 2.36 | 493V+501F+374Q+391S | 2.37 | 2.81 |
| N501F | 1.14 | 1.79 | 501Y+391S | 1.43 | 1.93 | 494K+501Y+391S | 1.81 | 2.25 | 493M+494K+501Y+391S | 2.34 | 2.73 |
| Q493M | 1.12 | 1.40 | 501F+518G | 1.41 | 2.03 | 453F+501Y+391S | 1.79 | 2.45 | 493M+494K+501F+391S | 2.33 | 2.83 |
| N501Y | 1.11 | 1.66 | 501F+391R | 1.38 | 1.99 | 453F+501F+391S | 1.76 | 2.52 | 493L+501F+374Q+391S | 2.31 | 2.69 |
| Q498H | 1.10 | 1.70 | 501F+391T | 1.38 | 2.00 | 478I+501F+391S | 1.75 | 2.88 | 493M+501F+374Q+391S | 2.28 | 2.80 |
| Q493L | 1.09 | 1.38 | 501F+391K | 1.37 | 1.92 | 453K+501Y+391S | 1.73 | 2.26 | 493M+494K+501Y+391T | 2.27 | 2.67 |
| L452K | 1.07 | 1.31 | 501F+518T | 1.37 | 2.08 | 494K+501Y+391T | 1.72 | 2.17 | 493M+494M+501Y+391S | 2.27 | 2.63 |
| Q493F | 1.05 | 1.24 | 501F+518S | 1.37 | 2.03 | 494K+501F+391T | 1.72 | 2.26 | 493L+494K+501Y+391S | 2.26 | 2.61 |
| Q493A | 1.03 | 1.31 | 501Y+518G | 1.36 | 1.89 | 453F+501Y+391T | 1.72 | 2.37 | 493M+494M+501F+391S | 2.26 | 2.73 |
| S494K | 1.03 | 1.25 | 501Y+391R | 1.35 | 1.85 | 494K+498H+391S | 1.71 | 2.12 | 493M+494K+501F+391T | 2.26 | 2.75 |
| Q498W | 1.02 | 1.26 | 493V+503R | 1.35 | 1.59 | 501F+519G+391S | 1.71 | 2.39 | 493V+501F+519G+391S | 2.25 | 2.86 |
| Q493Y | 1.02 | 1.32 | 498H+518T | 1.35 | 2.05 | 478I+501Y+391S | 1.71 | 2.70 | 493V+494M+501Y+391S | 2.25 | 2.59 |
| P527M | 1.01 | 1.18 | 501Y+518T | 1.34 | 1.96 | 494K+501Y+391R | 1.71 | 2.17 | 493L+494K+501F+391S | 2.25 | 2.70 |
| Q498Y | 1.01 | 1.44 | 501Y+391K | 1.34 | 1.77 | 494M+501F+391S | 1.71 | 2.21 | 493V+494K+501Y+391S | 2.24 | 2.58 |
| Q493I | 1.00 | 1.16 | 501F+517R | 1.34 | 1.99 | 501F+374Q+391S | 1.70 | 2.15 | 453F+493L+501Y+391S | 2.24 | 2.79 |
| L452Q | 1.00 | 1.17 | 501F+391N | 1.34 | 1.87 | 494K+501F+391R | 1.70 | 2.26 | 493V+494K+501F+391S | 2.24 | 2.69 |
| Y453F | 0.98 | 1.41 | 501F+517G | 1.34 | 1.98 | 494K+501F+518G | 1.70 | 2.22 | 493V+501F+519S+391S | 2.23 | 2.85 |
| L452S | 0.97 | 1.11 | 498H+391S | 1.34 | 1.83 | 501F+519S+391S | 1.70 | 2.38 | 493V+494M+501F+391S | 2.23 | 2.68 |
| S373R | 0.97 | 1.10 | 501Y+391T | 1.34 | 1.84 | 494M+501Y+391S | 1.70 | 2.11 | 493L+501F+519G+391S | 2.22 | 2.82 |
| L452R | 0.97 | 1.12 | 484K+501F | 1.34 | 2.14 | 494K+501Y+518G | 1.70 | 2.14 | 493M+494K+501Y+391R | 2.22 | 2.64 |

**Table S6.** Scanning results for chicken with up to 4 AA changes on SARS-CoV-2 RBD.

| 1-AA change | *K*_D,app_ ratio chick | *K*_D,app_ ratio human | 2-AA  change | *K*_D,app_ ratio chick | *K*_D,app_ ratio human | 3-AA  change | *K*_D,app_ ratio chick | *K*_D,app_ ratio human | 4-AA  change | *K*_D,app_ ratio chick | *K*_D,app_ ratio human |
| --- | --- | --- | --- | --- | --- | --- | --- | --- | --- | --- | --- |
| N501F | 1.10 | 1.79 | 439G+501Y | 1.35 | 1.91 | 439G+498H+501Y | 1.96 | 2.66 | 439G+494K+498H+501Y | 2.28 | 3.04 |
| N501Y | 1.09 | 1.66 | 440G+501Y | 1.32 | 1.96 | 440G+498H+501Y | 1.85 | 2.68 | 439G+494M+498H+501Y | 2.28 | 2.95 |
| N501W | 0.97 | 1.21 | 453F+501Y | 1.32 | 2.14 | 498H+501Y+358W | 1.85 | 2.80 | 439G+498H+501Y+384R | 2.22 | 2.87 |
| L452K | 0.91 | 1.31 | 501Y+358W | 1.31 | 2.13 | 448G+498H+501Y | 1.85 | 2.77 | 439G+498H+501Y+395W | 2.21 | 2.97 |
| Q493V | 0.89 | 1.34 | 450G+501Y | 1.30 | 1.97 | 498A+501Y+358W | 1.84 | 2.74 | 439G+498H+501Y+390W | 2.19 | 3.00 |
| E484K | 0.87 | 1.32 | 448G+501Y | 1.29 | 2.03 | 498H+501F+358W | 1.82 | 2.84 | 439G+498H+501Y+367Q | 2.19 | 2.90 |
| N501V | 0.85 | 1.27 | 501F+358W | 1.29 | 2.22 | 439G+484K+501Y | 1.82 | 2.29 | 439G+498H+501Y+339D | 2.18 | 2.95 |
| Q493M | 0.85 | 1.40 | 482K+501F | 1.29 | 1.83 | 498T+501Y+358W | 1.81 | 2.82 | 439G+494E+498H+501Y | 2.18 | 2.91 |
| Q498W | 0.85 | 1.26 | 482K+501Y | 1.28 | 1.75 | 453F+498H+501Y | 1.81 | 2.77 | 439G+498H+501Y+369W | 2.18 | 2.95 |
| P527M | 0.84 | 1.18 | 453F+501F | 1.28 | 2.17 | 439G+493M+501Y | 1.81 | 2.50 | 439G+498H+501Y+384K | 2.17 | 2.81 |
| L452Q | 0.84 | 1.17 | 501F+518T | 1.28 | 2.08 | 450G+498H+501Y | 1.81 | 2.63 | 439G+498H+501Y+517W | 2.17 | 2.96 |
| Q498H | 0.83 | 1.70 | 501F+518S | 1.27 | 2.03 | 498H+501Y+518T | 1.81 | 2.70 | 439G+455F+498H+501Y | 2.17 | 2.69 |
| S494K | 0.82 | 1.25 | 501F+367Q | 1.27 | 2.02 | 498H+501F+518T | 1.81 | 2.76 | 439G+498H+501Y+365W | 2.17 | 2.99 |
| N501M | 0.81 | 1.17 | 501F+518G | 1.27 | 2.03 | 498H+501F+518S | 1.78 | 2.70 | 439G+498H+501Y+367N | 2.16 | 2.89 |
| N501I | 0.81 | 1.05 | 501Y+518T | 1.26 | 1.96 | 439G+498Y+501Y | 1.78 | 2.30 | 439G+494F+498H+501Y | 2.16 | 2.79 |
| Y369W | 0.81 | 1.27 | 501Y+392W | 1.26 | 1.99 | 498H+501Y+518S | 1.78 | 2.63 | 439G+494H+498H+501Y | 2.16 | 2.78 |
| Y365W | 0.81 | 1.27 | 501Y+367Q | 1.26 | 1.90 | 498H+501Y+392W | 1.77 | 2.67 | 439G+482K+498H+501Y | 2.16 | 2.72 |
| L452R | 0.81 | 1.12 | 501F+339D | 1.26 | 2.05 | 498A+501Y+518T | 1.77 | 2.61 | 439G+498H+501Y+395Y | 2.15 | 2.89 |
| L452S | 0.81 | 1.11 | 482R+501F | 1.26 | 1.77 | 498A+501Y+392W | 1.77 | 2.61 | 439G+494D+498H+501Y | 2.15 | 2.88 |
| Q493A | 0.80 | 1.31 | 501Y+518S | 1.26 | 1.90 | 453F+498A+501Y | 1.77 | 2.69 | 439G+335K+498H+501Y | 2.15 | 2.87 |

**Table S7.** Hyperparameter of CNN_seq model subject to optimization with search spaces and final optimized values.

| **Hyperparameter** | **Search space** | **Optimized value** |
| --- | --- | --- |
| Kernel size (Conv1) | [3, 15] | 6 |
| Kernel size (Conv2) | [3, 15] | 12 |
| Number of output channels (Conv1) | [4, 64] | 48 |
| Number of neurons (FC1) | [64, 512] | 192 |
| Learning rate | [0.0001, 0.01] | 0.0002189 |
| Weight decay (Adam optimizer) | [0.0001, 0.01] | 0.0005564 |
| Dropout rate (FC1, FC2) | [0.2, 0.9] | 0.5045 |
